# Focused learning promotes continual task performance in humans

**DOI:** 10.1101/247460

**Authors:** Timo Flesch, Jan Balaguer, Ronald Dekker, Hamed Nili, Christopher Summerfield

## Abstract

Humans can learn to perform multiple tasks in succession over the lifespan (“continual” learning), whereas current machine learning systems fail. Here, we investigated the cognitive mechanisms that permit successful continual learning in humans. Unlike neural networks, humans that were trained on temporally autocorrelated task objectives (focussed training) learned to perform new tasks more effectively, and performed better on a later test involving randomly interleaved tasks. Analysis of error patterns suggested that focussed learning permitted the formation of factorised task representations that were protected from mutual interference. Furthermore, individuals with a strong prior tendency to represent the task space in a factorised manner enjoyed greater benefit of focussed over interleaved training. Building artificial agents that learn to factorise tasks appropriately may be a promising route to solving continual task performance in machine learning.

**Significance Statement:** Humans learn to perform many different tasks over the lifespan, such as speaking both French and Spanish. The brain has to represent task information without mutual interference. In machine learning, this "continual learning" is a major unsolved challenge. Here, we studied the patterns of errors made by humans and state-of-the-art deep networks whilst they learned new tasks from scratch and without instruction. Humans, but not machines, seem to benefit from training regimes that focussed on one task at a time, especially when they had a prior bias to represent stimuli in a way that facilitated task separation. Machines trained to exhibit the same prior bias suffered less interference between tasks, suggesting new avenues for solving continual learning in artificial systems.

## Introduction

Intelligent systems must learn to perform multiple distinct tasks over their lifetimes whilst avoiding mutual interference among them [1]. Building artificial systems that can exhibit this “continual” learning is currently an unsolved problem in machine learning research [2]. Despite achieving high levels of performance when training samples are drawn at random (“interleaved” training), standard supervised neural networks fail to learn continually in settings characteristic of the natural world, where one objective is pursued for an extended timeperiod before switching to another (“focussed” training). After first learning task A, relevant knowledge is overwritten as network parameters are optimised to meet the objectives of a second task B, so that the agent “catastrophically” forgets how to perform task A [3]. For example, state-of-the-art deep reinforcement learning systems can learn to play several individual Atari 2600 games at superhuman levels, but fail over successive games unless their network weights are randomly reinitialised before attempting each new problem [4, 5].

Human evolution, however, appears to have largely solved this problem. Healthy humans have little difficulty learning to classify a fixed stimulus set along multiple novel dimensions encountered in series. For example, we can effortlessly categorise an individual flexibly according to age, race or gender, and even if a conceptually novel axis of classification (e.g. social status) is introduced, it rarely corrupts or distorts past category knowledge. A longstanding theory argues that humans overcome this challenge via hippocampal-dependent learning mechanisms that interleave ongoing experiences with recalled memories of past training samples [6, 7]. This interleaving serves to decorrelate inputs in time, and avoids catastrophic interference by preventing the network from successively overfitting to each task in turn. Indeed, allowing neural networks to store and “replay” memories from an episodic buffer can accelerate training in otherwise autocorrelated environments such as video games [4, 8]. In psychology, a rich literature has explored the relative costs and benefits of focussed and interleaved training on category learning, with a majority of studies reporting an advantage for mixing exemplars from different categories, rather than focussing on one category at a time [9,10]. Humans also typically show an advantage for interleaved training during skilled motor performance, such as in sports [11] or language pronounciation [12], and even in the acquisition of abstract knowledge, such as mathematical concepts [13]. These findings support a single, general theory that emphasises the benefits of interleaved training for continual task performance in humans and neural networks.

Here, however, we desribe an experiment that suggests a different view. We compared the mechanisms that promote continual task performance in humans and neural networks, focussing on a canonical cognitive paradigm, that involves switching between classification tasks with orthogonal rules. Whilst much is known about the factors that limit task switching performance in explicitly-instructed, rule-based paradigms [14], here we explored how humans learned to switch between tasks from scratch and without prior instruction. In other words, rather than studying the control processes that permit task switching, we investigated how a general problem composed of multiple subtasks is learned by trial and error alone. We taught human and artificial agents to classify high-dimensional, naturalistic stimuli (trees) according to whether they were more or less leafy (task A) or more or less branchy (task B), drawing trial-unique exemplars from a uniform bidimensional space of leafiness and branchiness (**Fig.1a**).

**Figure 1.**
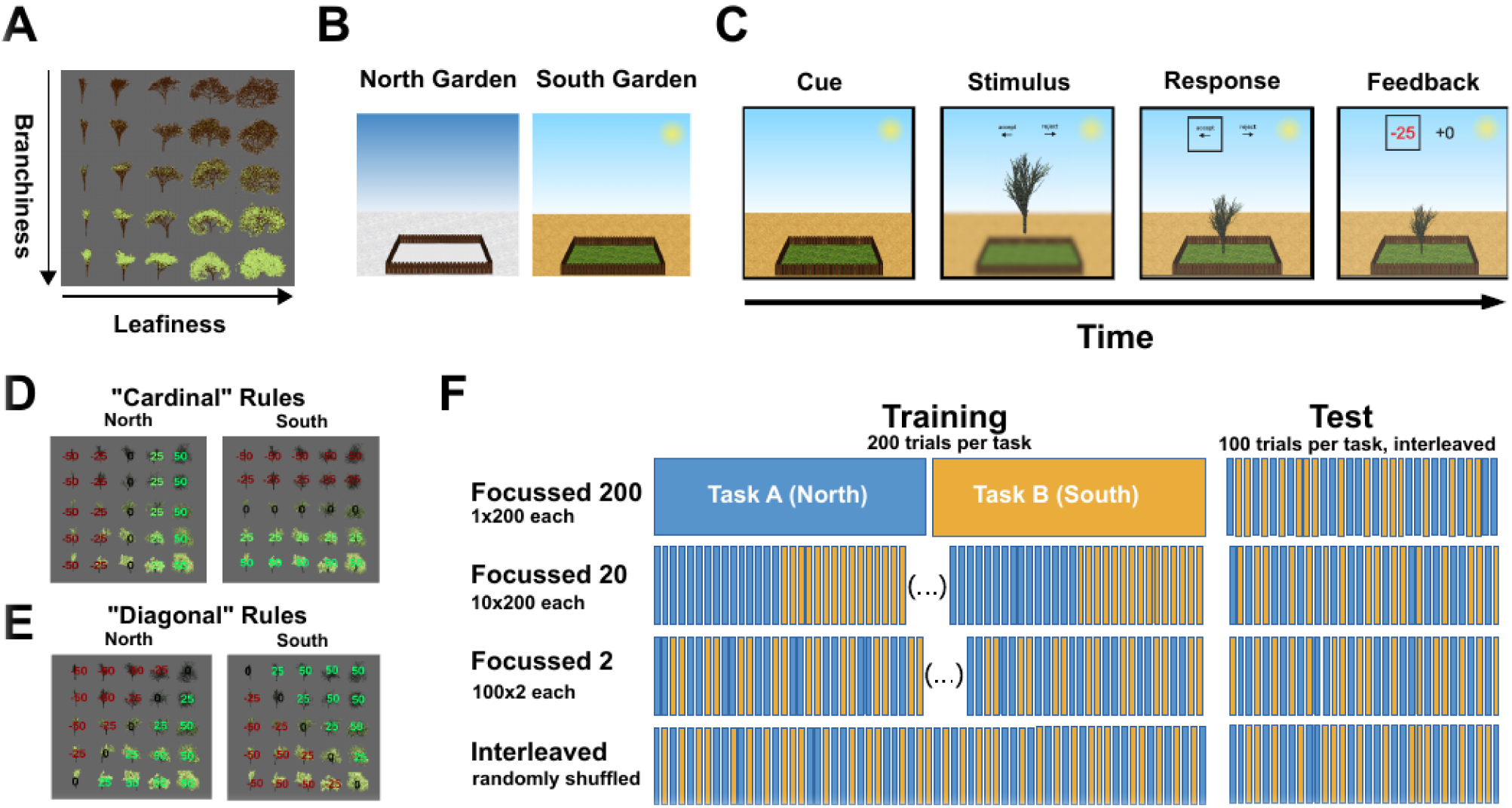
Task Design, Experiment 1. (**A**) Naturalistic tree stimuli were parametrically varied along two dimensions (leafiness and branchiness). (B) All participants enganged in a virtual gardening task with two different gardens (north and south). Via trial and error, they had to learn which type of tree grows best in each garden. (C) Each training trial consisted of a cue, stimulus, response and feedbak period. At the beginning of each trial, an image of one of the two gardens served as contextual cue. Next, the context was blurred and the stimulus (tree) appeared together with a reminder of the key mapping (“accept” vs “reject”, corresponding to “plant” vs “don’t plant”) in the center of the screen. Once the participant had communicated her decision via button press (left or right arrow key), the tree would either be planted inside the garden (“accept”) or disappear (“reject”). In the feedback period, the received and counterfactual rewards were displayed above the tree, with the received one being highlighted, and the tree would either grow or shrink, proportionally to the received reward. Test trials had the same structure, but no feedback was provided. Key mappings were counterbalanced across participants (D) Unbeknownst to the participants a priori, there were clear mappings of feature dimensions onto rewards. In Exp 1a (cardinal group), each of the two feature dimensions (branchiness or leafiness) was mapped onto one task rule (north or south). The sign of the rewards was counterbalanced across participants. (E) In Exp 1b (diagonal group), feature *combinations* were mapped onto rewards, yielding nonverbalisable rules. Once again, we counterbalanced the sign of the rewards across participants (F) Exp 1a and 1b were between-group designs. All four groups were trained on 400 trials (200 per task) and evaluated on 200 trials (100 per task). The groups different in the temporal autocorrelation of the tasks during training, ranging from “focussed 200” (200 trials of one task, thus only one switch) to “interleaved” (randomly shuffled and thus unpredictable task switches). Importantly, all four groups were evaluated on interleaved test trials. The order of tasks for the focussed groups was counterbalanced across participants.

We found that relative to interleaved training, providing humans (but not neural networks) with autocorrelated objectives promoted learning and protected task representations from mutual interference, even on a later generalisation test in which the two tasks were conducted on new stimuli that were interleaved over trials. Fitting a psychophysical model to the data, we found that focussed learning promoted accurate representations of two orthogonal decision boundaries required to perform the task (i.e. to “factorise” the problem into two discrete tasks), whereas interleaved training encouraged humans to form a single linear boundary that failed to properly disentangle the two tasks. This benefit was greatest for those individuals whose prior representation of the stimulus space (as measured by pre-experimental similarity judgments among exemplars) organised the stimuli along the cardinal task axes of leafiness and branchiness.

Suprisingly, we even found evidence for the protective effect of focussed learning after rotating the category boundaries such that rules were no longer verbalisable. These findings suggest that autocorrelated training objectives encourage humans (but not current neural networks) to factorise complex tasks into orthogonal subcomponents that can be represented without mutual interference.

## Results

In **Exp.1a** adult humans (n ~ 200) performed a “virtual gardening task” that required them to learn to plant naturalistic tree stimuli in different gardens (north vs. south, denoted by different images) (**Fig. 1b,c**). Unbeknownst to participants, different features of the trees (leafiness vs. branchiness) predicted growth success (and thus reward) in either garden (**Fig. 1d**). Different cohorts learned to plant (classify) trees under training regimes in which gardens (tasks) switched randomly from trial to trial (interleaved group), remained constant over sequences of 200 trials (F200 group), 20 trials (F20 group), or 2 trials (F2 group) (**Fig. 1f**). All learning was guided by trialwise feedback alone; we were careful not to alert participants either to the rules or to the cardinal dimensions of the tree space (leafiness, branchiness). Our critical dependent measure was performance on a final generalisation test session involving novel tree exemplars in which leafy and branchy tasks were interleaved but no feedback was provided.

We first describe three observations that we found surprising. Firstly, the F200 group (which received the most focussed training) performed overall best during interleaved test (ANOVA: *F_3,172_* = 5.06, *p* < 0.05; F200 > Interleaved: *t_93_* = 2.324, *p* < 0.05; F200 > F2: *t_86_* = 3.814, *p* < 0.001) (**Fig. 2a,b**). This is striking, given the encoding specificity benefit that the rival interleaved group should have received from the shared context during learning and evaluation [15]. Secondly, the benefits of focussed learning were observed even when analysis was limited to switch trials at test (ANOVA: *F_3,172_* = 4.59, *p* < 0.01; F200 > Interleaved: *t_93_* = 2.062, *p* < 0.05; F200 > F2: *t_86_* = 3.590, *p* < 0.001) despite the fact that participants in the F200 had only experienced a single switch during training (**Fig. 2c**). We found this remarkable, given that training on task switching has previously been shown to reduce switch costs [16]; our data suggest that task switching can be paradoxically facilitated without any switch practice. Finally, this benefit did not merely arise from the heightened predictability of the training sequence under focussed learning: the F2 group, which experienced an identical number of task switches during training as the interleaved group but in predictable order, performed worst overall at test (one-sided unpaired t-test. F2 < Interleaved: *t_85_* = 1.834, *p* < 0.05; **Fig. 2b**).

**Figure 2.**
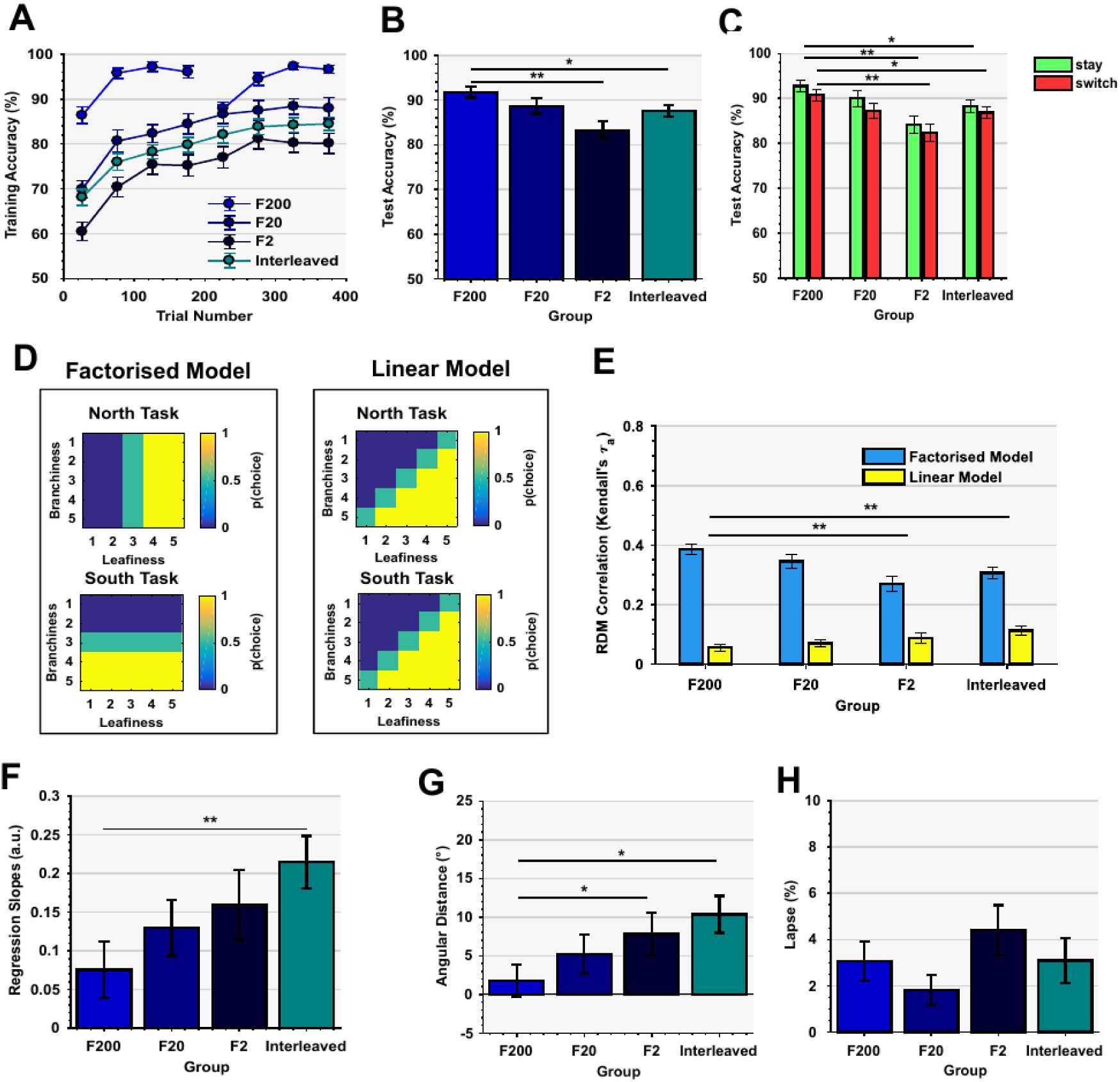
Results of Experiment 1a. (A) Training curves, depicting mean training accuracy, averaged over 50 trials. All four groups reached a stable performance plateau by the end of the training phase. Both, the learning rate and the final training performance depended on the degree of temporal autocorrelation of the tasks during training. Error bars depict SEM. (B) The mean performance on new tree stimuli during the test phase differed significantly between groups, as the F200 group performed significantly better than the F2 and Interleaved groups. Error bars depict SEM. (C) Mean Test performance separately for switch and stay trials. Surprisingly, even on switch trials, the F200 group outperformed the interleaved and F2 group, despite having experienced only one task switch during training. Error bars depict SEM. (D) Conceptual choice probability models. The factorised model (left) predicted that participants learned two separate boundaries for each task, corresponding to the rewards that were assigned to each dimension in trees space. The linear model (right) was based on the assumption that participants had learned the same, *linear* boundary for both tasks, separating the trees space roughly into two halfes that yielded equal rewards and penalties in both tasks. (E) Results of RDM model correlations on test phase data. Whilst the factorised model provided a better fit to the data for all groups, its benefit over the linear model was greatest for the F200 than for the F2 and Interleaved group. Error bars depict SEM. (F) Slope of Sigmoids fit to the test-phase choice proportions of the task-irrelevant dimension. The significantly steeper slope for interleaved compared to F200 indicates stronger *intrusion* from the task-irrelevant dimension. Error bars depict SEM (G) Mean angular distances between true category-boundary and single-subject decision boundary, estimated by the psychophysical model. A significantly stronger bias for F2 and Interleaved compared to F200 suggests that focussed training allows precise recall of the decision boundary, even during an interleaved test phase. Error bars depict SEM. (H) Mean lapse rates, obtained from the same psychophysical model. There were no significant differences between groups. Error bars are SEM.

We next explored how focussed learning promoted task performance in humans using three more detailed analysis strategies. First, we plotted psychometric curves that revealed how choices at test depended on both the relevant dimension (e.g. leafiness) and the irrelevant dimension (e.g. branchiness). Comparing the slopes of these functions revealed that focussed learning reduced the impact of the irrelevant dimension on choice, as if F200 training prevented intrusions from the rival task (F200 < Interleaved: *Z* = 2.966, *p* < 0.01; **Fig. 2f**). Secondly, we plotted choice probabilities p(plant) at test for each tree across the bidimensional space, to visualise the decision boundary learned in each training regime. Visual inspection suggested that whereas F200 training allowed participants to learn two orthogonal decision boundaries that cleanly segregated trees in leafiness and branchiness conditions, after interleaved training participants were more prone to use a single, diagonal boundary through 2D tree space that failed two disentangle the two tasks (**Fig. S1a**). Using an approach related to representational similarity analysis (RSA) [17], we correlated human choices with those predicted by models that learned the best possible linear boundary in tree space (linear model) or two boundaries that cleaved different compressions of the tree space optimally according to the relevant dimension (factorised model; **Fig. 2d**). Although the factorised model fit human data better than the linear model in all conditions, its advantage was greatest in the F200 condition (F200_tau_diff_ > Int_tau_diff_: *Z* = 3.592, *p* < 0.001; F200_tau_diff_ > F2_tau_diff_: *Z* = 3.699, *p* < 0.001; **Fig 2e**). Thirdly, we combined elements of these analysis approaches, fitting a more flexible psychophysical model to the data that estimated both the most likely angle of the bound and the noise and fraction of random errors (lapses) in sensory judgment. The fits revealed that the benefit in the F200 condition was due to both a sharper and more accurate bound (F200 < Int: *Z* = 2.564, *p* < 0.05; F200 < F2: *Z* = 2.024, *p* < 0.05; **Fig. 2g**), but not a different lapse rate (Kruskal-Wallis: H_3,172_ = 3.87, *p* = 0.275; **Fig. 2h**). Together, these findings suggest that focussed training helps promote two robust representations of the two tasks from which the problem was composed (‘factorised learning’). This learning strategy facilitates performance when tasks are later encountered in random succession.

Categorisation can rely on explicit rule discovery or more implicit learning of stimulus-response associations [18]. In order to test whether the benefit of focussed learning was due to the promotion of rule-based strategies, next we rotated the category boundaries for tasks A and B by 45º in tree space such that they lay along a non-verbalisable leafy/branchy axis (‘diagonal boundary’ condition), and repeated our experiment in a new cohort of participants (Exp.1b; n = 200). Interestingly, although we observed no significant differences in test performance among the different training groups (ANOVA on accuracy: *F_3,162_* = 0.25, *p* = 0.863; stay trials only: *F_3,162_* = 0.37, *p* = 0.776; switch trials only: *F_3,162_* = 0.63, *p* = 0.598) (**Fig. 3a,b**), we still saw evidence of factorised learning in the F200 condition. This was demonstrated by shallower psychometric slopes for the irrelevant dimension at test (F200 < Int: *Z* = 3.106, *p* < 0.001; F200 < F2 *Z* = 2.899, *p* < 0.01; **Fig 3f**), and a better fit to RDMs for the factorised task model in the F200 condition (F200 > Int: *Z* = 3.271, *p* < 0.001; F200 > F2: *Z* = 3.57, *p* < 0.001; **Fig 3e**), both relative to interleaved and F2 training. Indeed, using the psychophysical model we observed lower estimates of boundary error in the F200 condition (F200 < Int *Z* = 3.209, *p* < 0.001; F200 < F2 *Z* = 2.695, *p* < 0.01; **Fig 3g**), but higher lapse rates (F200 > Int *Z* = 1.965, *p* < 0.05; F200 > F2 *Z* = 1.739, *p* < 0.05; **Fig 3h**). Thus, it seems that under nonverbalisable boundaries, focussed training promotes learning of the effective boundary, but this benefit only offsets (but does not reverse) a nonspecific cost that may have been incurred by limited experience with task switching during training.

**Figure 3.**
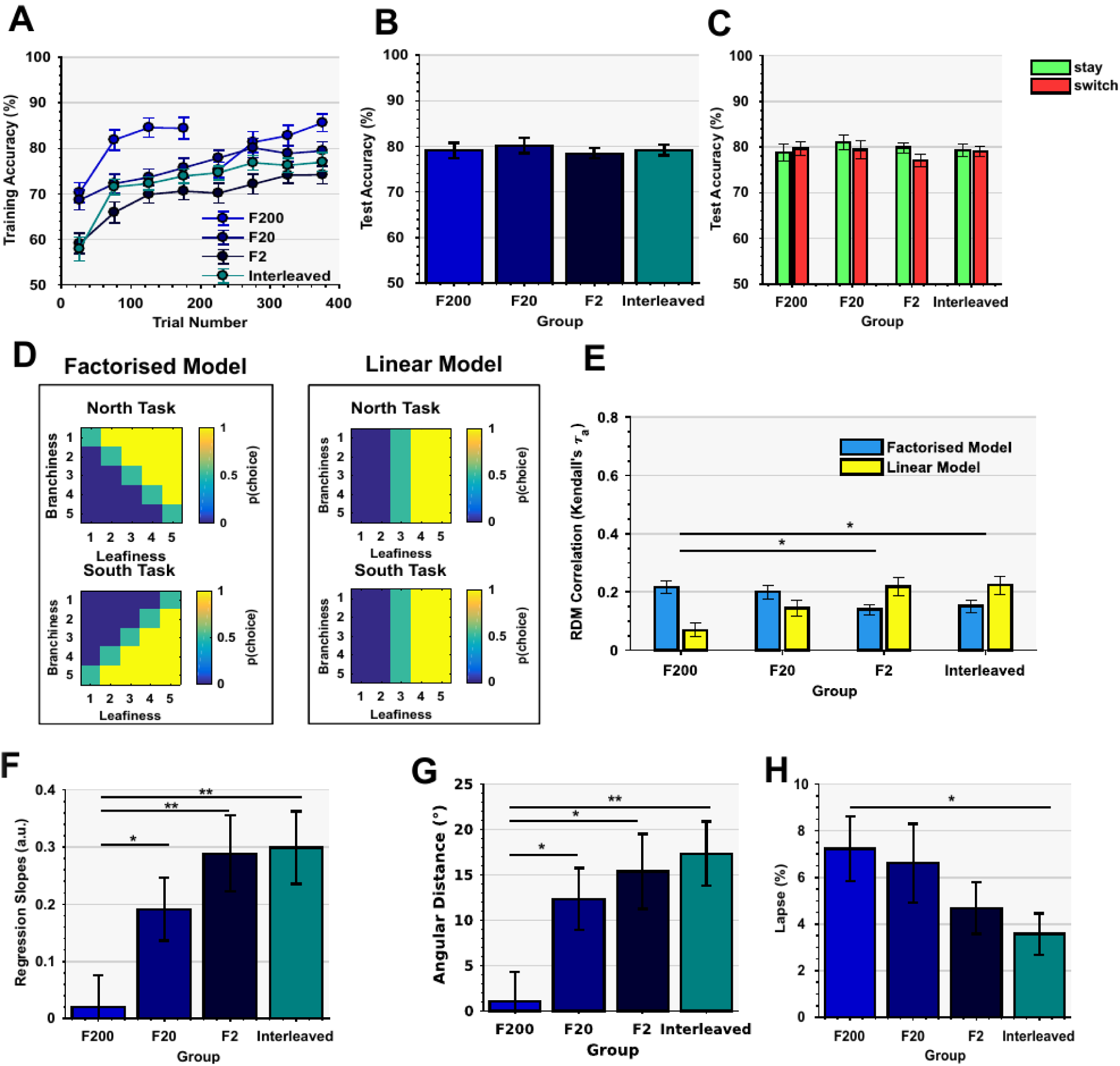
Results of Experiment 1b. (A) Training curves. At the end of the training session, all four groups had reached a stable performance plateau. Error bars depict SEM. (B) Mean test performance. In contrast to Exp1a, there was no significant difference in performance between groups. Error bars depict SEM. (C) Performs did also perform equally well on stay and switch trials. Error bars depict SEM. (D) Conceptual model RDMs. The same reasoning applies as described in. (E) RDM model correlations at test. Despite equal test performance, the relative advantage of the F200 over the linear model is stronger for F200 than for F2 or interleaved, suggesting that focussed training did result in better task-separation, despite equal performance. Error bars depict SEM. (F) Slope of sigmoidal fits to choice probabilities to feature values in the task-irrelevant dimension. Intrusion was weakest for F200 and larges for F2 and Interleaved groups. Error bars depict SEM. (G) Mean bias of the decision boundary obtained by the psychophysical model. The bias was smallest for F200, indicating that focussed training did allow the participants to form precise estimates of the category boundaries, in contrast to interleaved training. Error bars depict SEM. (H) Estimated mean lapse rates. The F200 group made a higher number of unspecific random errors during the test phase, compared to the Interleaved group, which explains equal test performance despite evidence for successful task factorisation. Error bars depict SEM.

One possibility is that focussed training allows humans to compress their potentially high-dimensional representation of tree space onto a single discrimination axis, where they can selectively apply the relevant linear boundary (e.g. leafy vs. branchy) according to the task context. If so, it follows that participants who are *a priori* predisposed to represent the trees in an orderly space of leafiness vs. branchiness might be most able to benefit from such a strategy, and thus from focussed training, especially when the boundaries lie on the cardinal axes of this space. Subsequently, thus, we repeated our experiments on a new participant cohort who classified the stimuli according to cardinal (**Exp.2a**) and diagonal (**Exp.2b**) boundaries, but this time we measured participants’ representation of tree space using a pre- and post-experimental “arena” task, in which trees were manually arranged according to their similarity or dissimilarity within a circular aperture. This allowed us to estimate the extent to which the prestimulus arrangement matched the 2D leafy vs. branchy grid encountered in the subsequent classification task(s), and quantify participants’ prior on the stimulus space being organised as a leafy x branchy grid (**Fig. 4**).

**Figure 4.**
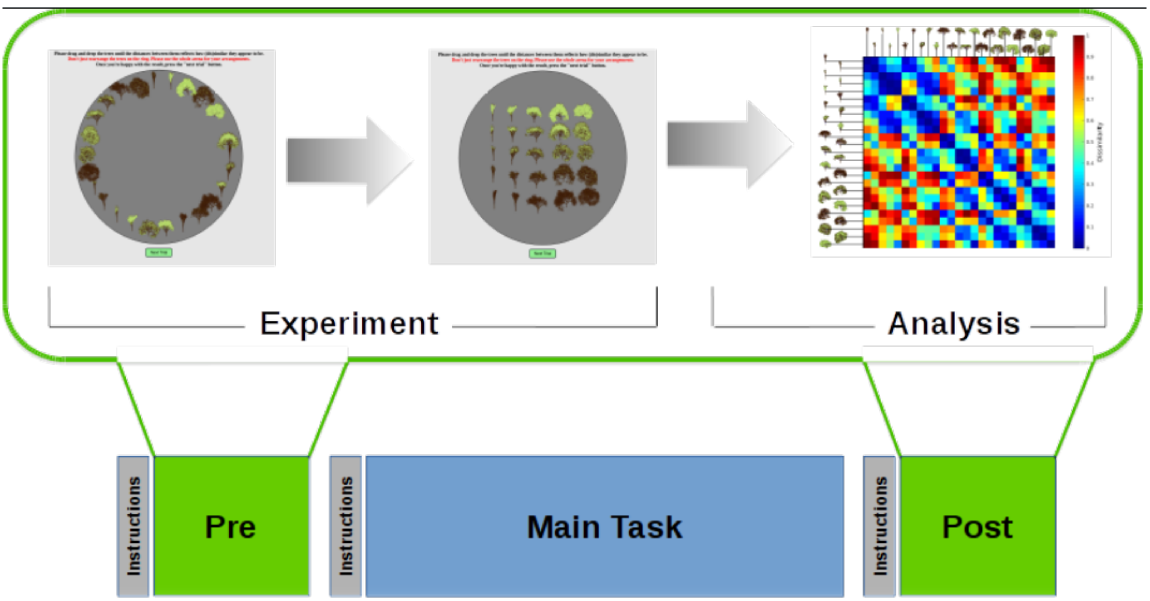
Task Design, Experiment 2. We tested four new cohorts on cardinal and diagonal boundaries and F200 vs Interleaved training regimes. Before and after the main experiment, which was identical to the procedures described in, participants engaged in an arena task, in which they were asked to rearrange 25 trees (one for each position in the 5x5 grid depicted in Fig. 1a) via mouse drag and drop inside a circular aperture until pairwise euclidean distances reflected subjective dissimilarity. During data analysis, we compute the single-trial normalised RDM, averaged the RDMs across trials, yielding one RDM per subject and phase. Correlation of the RDM obtained before subjects engaged in the task with a model RDM (which assumed perfect grid-like arrangement and thus awareness of the true 5x5 space) yielded a “grid” prior (Kendall tau) for each participant.

We then tested how this “grid” prior interacted with training regime to predict subsequent test performance. The results of the main task replicated those of Exp.1a and Exp.1b, with stronger evidence for factorised learning under F200 than interleaved training, as indicated by fits to choice matrices (Exp 2a: F200 > Int: *Z* = 3.213, *p* < 0.001; Exp 2b: F200 > Int *Z* = 2.310, *p* < 0.05), and estimates of boundary error (Exp 2a: F200 < Int: *Z* = 2.621, *p* < 0.01; Exp 2b: F200 < Int *Z* = 2.199, *p* < 0.05) from the psychophysical model (**Fig. S2**). However, of primary interest for this experiment was how participants’ prior representation of the stimulus space interacted with training regime to promote learning. We used the arena task to compute a single quantity (“grid prior”) that indicated the extent to which participants arranged the trees into a 2D grid defined by leafiness and branchiness in the pre-experimental session. Splitting participants according to their grid prior, in Exp. 2a (cardinal boundaries) we found that those who tended to represent the space a priori as a 2D grid of leafiness vs. branchiness exhibited more benefit from F200 training than those who did not (F200_highPrior_ > F200_lowPrior_: *Z* = 2.871, *p* < 0.005; Int_highPrior_ = Int_lowPrior_: *Z* = 1.692, *p* = 0.091; **Fig 5a**). The finding remained significant when we used an ANCOVA to estimate interactions between grid prior and training on performance (Grid prior: *F_1,134_* = 13.54, *p* < 0.001; Group: *F_1,134_* = 6.6, *p* < 0.05; Grid prior*Group: *F_1,134_* = 6.28, *p* < 0.05). Moreover, signatures of factorised learning, including the fits of the factorised model to human choice matrices, were more pronounced for the high grid prior group under F200 but not interleaved training (F200_highPrior_ > F200_lowPrior_: *Z* = 3.122, *p* < 0.01; Int_highPrior_ > Int_lowPrior_: *Z* = 1.55, *p* = 0.061; ANCOVA: Grid prior: *F1,134* = 22.36, *p* < 0.0001; Group: *F_1,134_* = 12.18, *p* < 0.001; Grid prior*Group: *F_1,134_* = 8.16, p < 0.01; **Fig. 5b**) By contrast, in the diagonal boundary case (Exp.2b), we observed generally higher performance and task factorisation for those participants who had a higher grid prior, but no interaction with training regime (Accuracy: highPrior_pooled_ > lowPrior_pooled_, *Z* = 2.052, *p* < 0.05; Factorisation: highPrior_pooled_ > lowPrior_pooled_, Z = 1.969, p < 0.05; ANCOVA on accuracy: Grid prior: *F_1,99_* = 5.96, *p* < 0.02, Group: *F_1,99_* = 1.5, *p* = 0.223, Grid prior*Group: *F_1,99_* = 0.31, *p* = 0.579; ANCOVA on correlations with factorised model: Grid prior: *F_1,99_* = 9.15, *p* < 0.01; Group: *F_1,99_* = 2.07, *p* = 0.153; Grid prior*Group: *F_1,99_* = 0.01, *p* = 0.919; **Fig 5c,d**).

**Figure 5.**
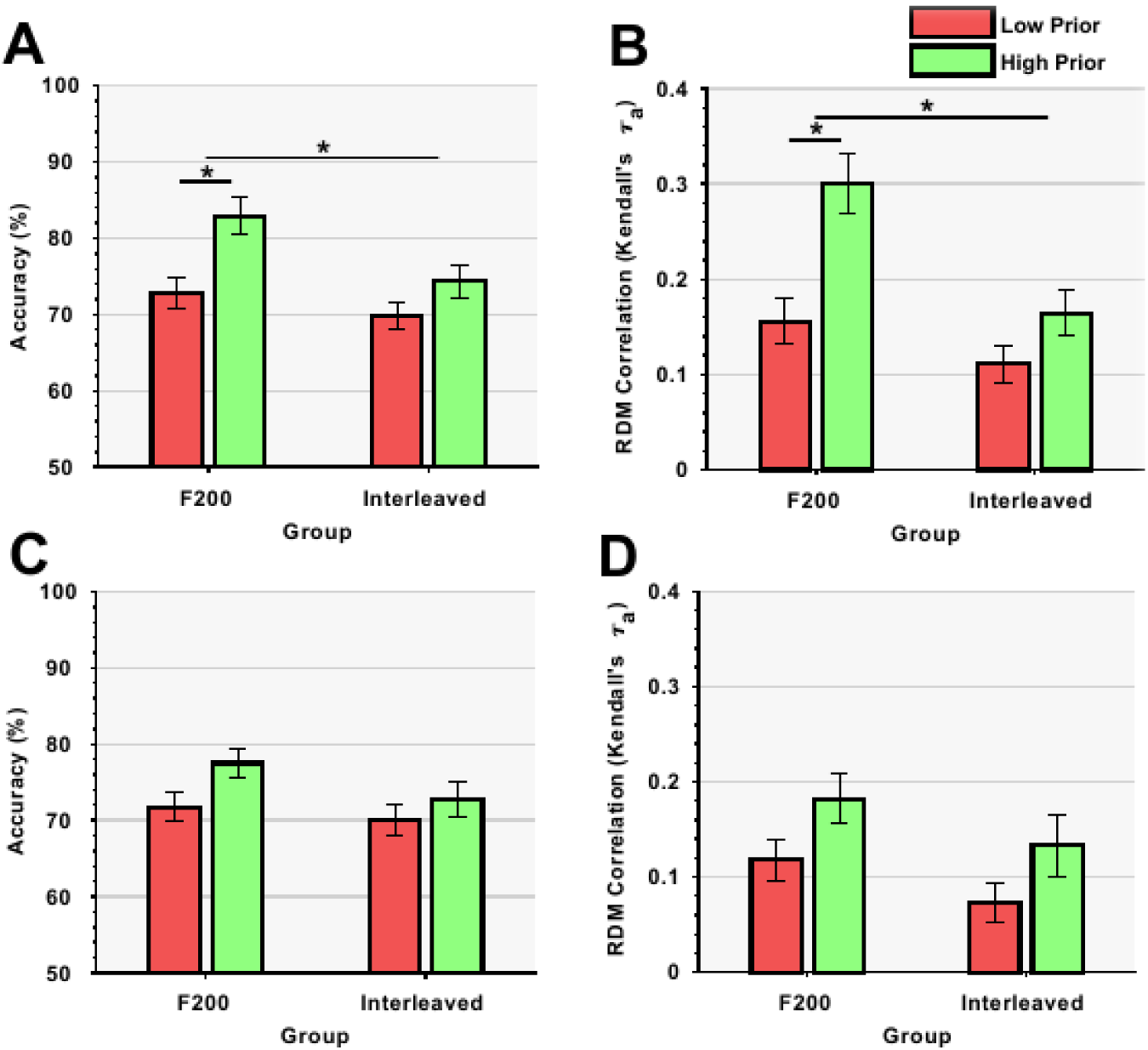
Results of Experiment 2. (A) Exp 2a (cardinal boundary): Median split of test performance. The benefit of focussed training was significantly stronger for participants with a higher prior on the structure of the trees space. Error bars depict SEM. (B) Exp 2a:Mediansplitofcorrelations between choice probabilities and factorized model().Under focussed training, articipants with a strong prior showed significantly stronger evidence of task factorisation. (C) Exp 2b (diagonal boundary). There was no difference between low and high grid priors on mean test accuracy. (D) Exp 2b. The correlation coefficients of the factorised model did not differ between groups. An ANCOVA (see main text) revealed a main effect of the prior on task factorisation, but no interaction with group.

Next, for comparison with humans, we trained neural networks to perform the task. In **Exp.3**, convolutional neural networks (CNNs) were separately trained on the cardinal and diagonal tasks under either focussed or interleaved training conditions, and classification performance was periodically evaluated with a held-out test set for which no supervision was administered. The networks received the image pixel values as inputs and were optimised (during training) with a supervision signal designed to match the reward given to humans. As expected, under interleaved training the network rapidly achieved ceiling performance at test on both tasks. However, under focussed learning conditions network performance dropped to chance after each task switch (**Fig. 6a,b**), in line with an extensive literature indicating that such networks suffer catastrophic interference in autocorrelated training environments [3,19]. Using an RSA approach, we correlated the layer-wise activity patterns with model Representational Dissimilarity Matrices (RDMs) that either assumed pixel value encoding (pixel model), category encoding of both tasks (factorised model), category encoding of the most recent task only (interference model) or a linear boundary through feature space (linear model). Unit activation similiarity patterns in the early layers reflected the pixelwise similarity among input images, whereas deeper layers exhibited a representational geometry that flipped with each new task, as predicted by the interference model, and indicative of catastrophic forgetting (**Fig. 6c,d**).

**Figure 6.**
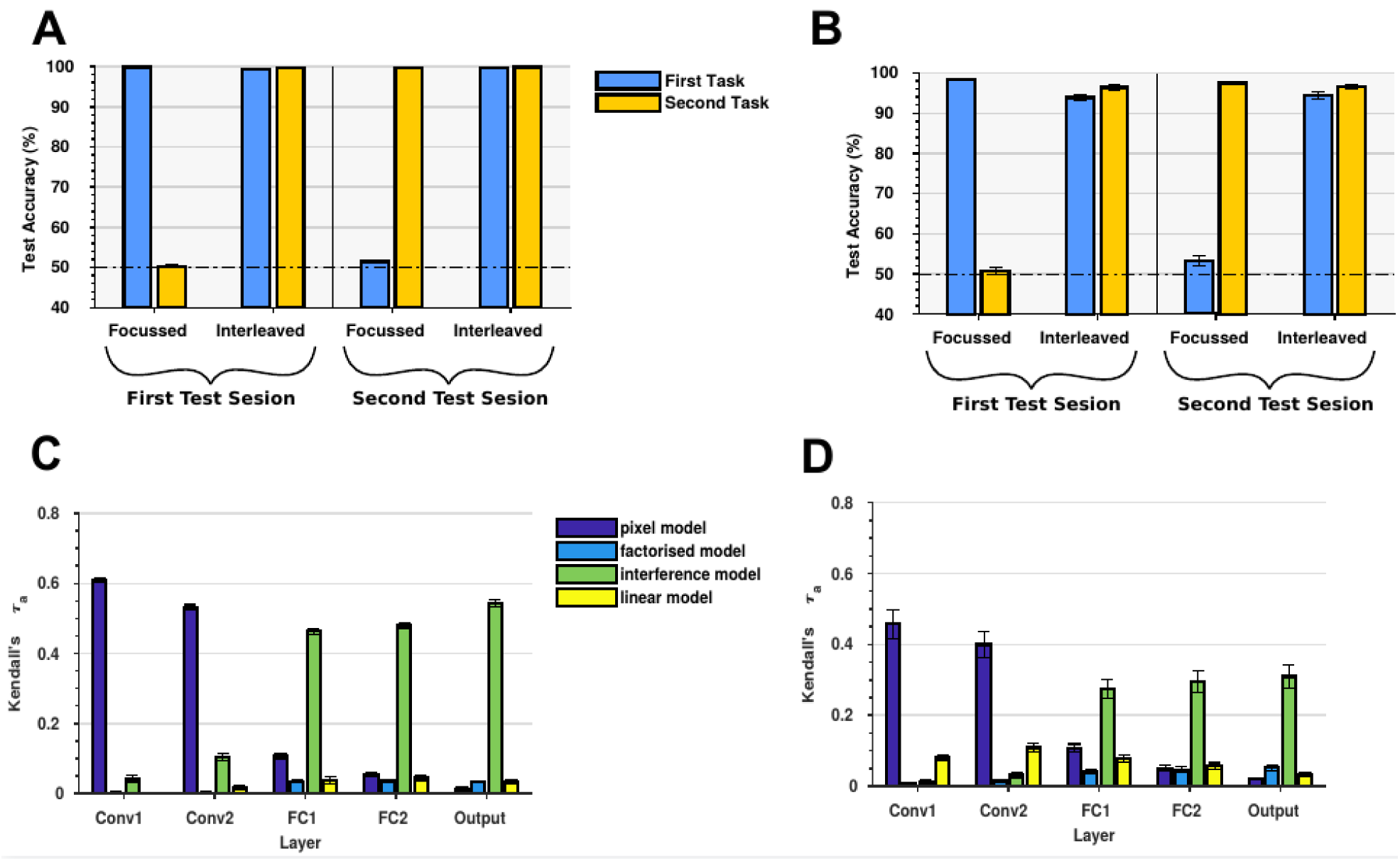
Results of Experiment 3. (A) Exp 3a (cardinal boundary). Mean performance of the CNN on independent test data,calculated after the first and second half of training, separately for the first and second task and focussed vs interleaved training. As expected, interleaved training yielded equal performance on both tasks which quickly reached ceiling. In contrast, the network trained with a focussed regime performed at ceiling for the first task, but dropped back to chance level after it had been trained on the second task, on which it did also achieve ceiling performance. Error bars depict SEM across independent runs. (B) Exp 3b (diagonal boundary) Mean test performance. We observed similar patterns as for the cardinal boundary: Focussed training resulted in catastrophic interference, whereas interleaved training allowed the network to learn both tasks equally well. Interestingly, the CNNs performed slightly worse on the diagonal boundary, similar to our human participants. Error bars depict SEM. (C) Exp 2a, focussed training. Layer-wise RDM correlations between RDMs optained from activity patterns and model RDMs. The correlation with the pixel dissimilarity model decreases from the first to last layer, whereas the correlation with a model which assumes catastrophic interference increases. Neither the factorised nor the linear model explain the data well, indicating that focussed training did not result in task factorisation or convergence towards a boundary which yields good performance in both contexts. Error bars indicate SEM. (D) Exp 2b, focussed training. We see similar patterns as for the model which was trained on a cardinal boundarty (): Correlations with the pixel model decrease and correlations with the interference model increase from the first convolutional to the output layer. Error bars depict SEM.

Our investigations of human task learning suggest that prior knowledge of the structure of the stimulus space is one factor that allows humans to learn task representations that are protected from mutual interference. We thus wondered whether pretraining the CNN to factorise the task space appropriately would mitigate the effect of catastrophic interference. To achieve this, we trained a deep generative model (a beta variational autoencoder or β-VAE [20]) on a large dataset of trees drawn from the 2D leafy x branchy space (**Exp 4a**). The autoencoder learned to reconstruct its inputs after passing signals through two “bottleneck” nodes, which obliges the network to form a compressed representation of the latent stimulus space. Unlike a standard VAE, which typically learns an unstructured latent representation, the β-VAE can learn disentangled and interpretable data generating factors, similar to a child which aquires structured visual concepts through passive observation of its surroundings [20]. To verify that the network had learned appropriately, we traversed the latent space of activations in the two bottleneck units and visualised the resulting tree reconstructions, revealing a 2D embedding space that was organised by leafiness x branchiness (**Fig. S3**).

Next, in **Exp.4b**, we retrained the CNN, but using the β-VAE encoder as a feature extractor for the main task. We provided the output of the encoder from the β-VAE an input to the first fully-connected layer in the CNN, thereby allowing it to utilise similar prior knowledge about the structure of the inputs, as our human participants seemed to do. This approach mirrors that which occurs during human development, in which rich knowledge of the statisical structure of the world is learned before the rules guiding behaviour explicitly taught [21].

The effect of catastrophic interference in the CNN, although still present, was reduced by this intervention. In particular, the network retained some knowledge of previous tasks it had experienced during focussed training (**Fig. 7a,d**), leading to overall improved performance in this condition relative to the vanilla CNN for the cardinal, but not the diagonal group (test accuracy across both tasks, focussed training: cardinal priorCNN > vanillaCNN Z = 4.972, p < 0.001, diagonal priorCNN = vanillaCNN Z = 1.43, p = 0.151). Using RSA to investigate representational geometry in the network, we found that using the autoencoder as a feature extractor encouraged the network to represent the task in a factorised manner, with RDMs showing reduced correlations with the interference model and increased correlations with the factorised model in both fully-connected layers (correlation with factorised model for priorCNN > vanillaCNN, all p-values < 0.001 for both FC layers and output layer; correlation with interference model for vanillaCNN > prior CNN, all p-values < 0.001; **Fig. 7c,d**). Interestingly, in the diagonal boundary case, the advantage for the factorised model was only significant in the output layer (Z = 2.19, p < 0.05; **Fig. 7e**), but we observed reduced correlations with the interference model in all three layers (all p-values < 0.001; **Fig. 7f**). We take these data as a proof-of-concept that unsupervised learning of the statistical structure of the world may be one important factor for promoting continual task performance in both humans and neural networks.

**Figure 7.**
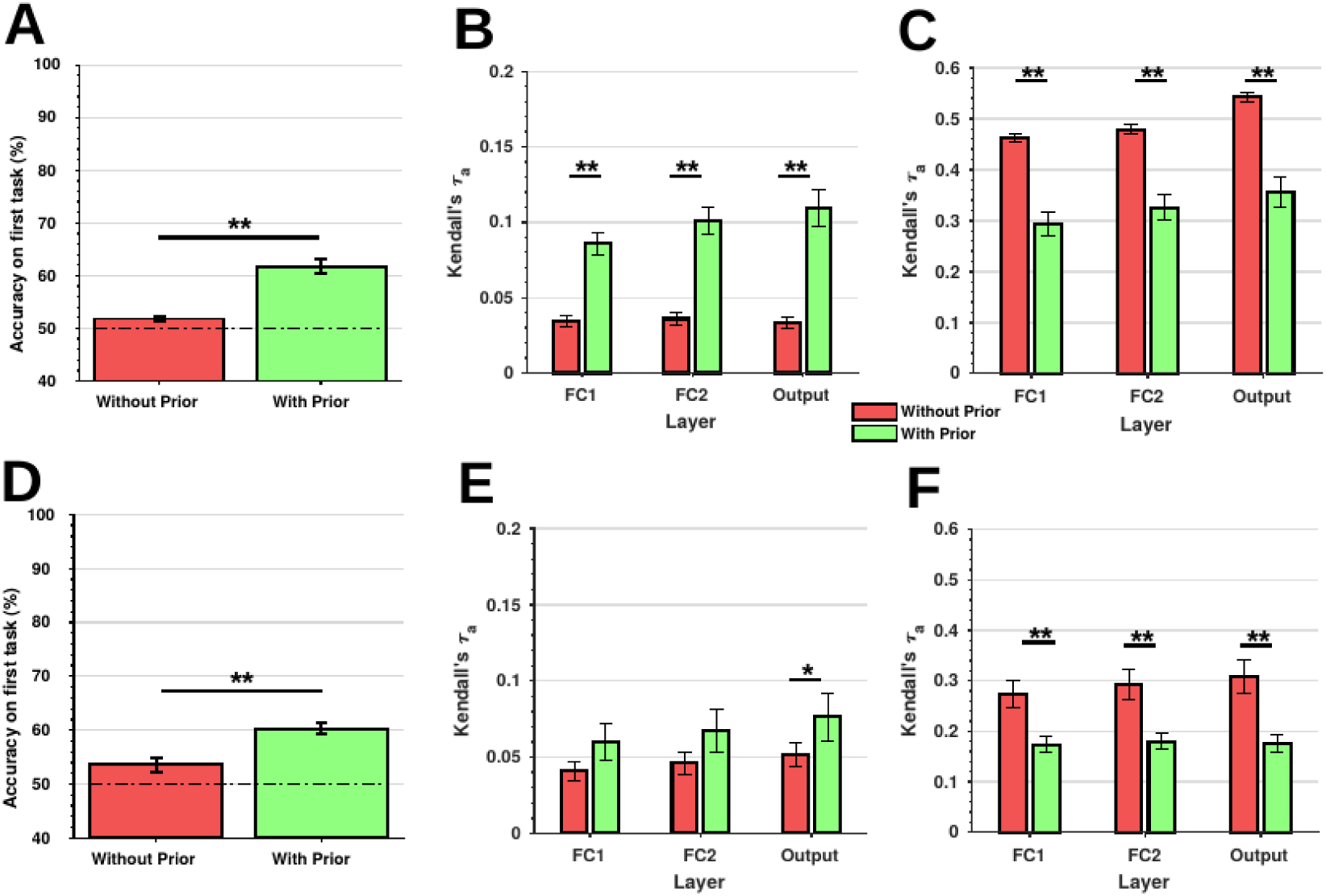
Results of Experiment 4. (A) Exp. 7a (cardinal), focussed training. Comparison of performance on the first task aftertraining on the second task, between the model from Exp 3 (“vanilla” CNN, without priors) and the model from Exp 4 (“pretrained” CNN, with priors from β-VAE encoder). Unsupervised pretraining of the data structure partially mitigated catastrophic interference. (B) Exp 7a, focussed training. Comparison of layer-wise RDM correlations with factorised model for CNN, between networks without and with unsupervised pretraining. The β-VAE encoder resulted in stronger correlations with the factorised model in each layer that was training on the two tasks. (C) Exp 7a, focussed training. Comparison of layer-wise RDM correlations with interference model. Likewise, pretraining significantly reduced correlations with the Catastrophic Interference model in each layer. (D) Exp 7b (diagonal), focussed training. Mean accuracy on the first task after training on the second task, for vanilla and pretrained CNN. Interestingly, pretraining did reduce the severity of interference also when the network was trained on diagonal boundaries. (E) Exp 7b, focussed training. RDM correlations with factorised model. If trained on a diagonal boundary, correlations with a factorised model increased only in the output layer. (F) Exp 7b, focussed training. RDM correlations with the interference model, however, increased significantly in each layer, even if the network was trained on a diagonal boundary. All error bars depict SEM.

## Discussion

Humans can continually learn to perform new tasks across the lifespan, but the computational mechanisms that permit continual learning are unknown. Here, we tackled the problem of understanding how humans learn two distinct classification tasks from scratch and without instruction. We first show that humans benefit from focussed learning conditions, in which task objectives are presented in an autocorrelated fashion, even when later test stimuli are interleaved over trials, a scenario that was not experienced during initial training. This result poses a challenge for standard computational models of learning, including current neural network accounts, where test performance is typically heavily determined by the level of exposure to equivalent training examples [15,22].

Secondly, our behavioural anlaysis offers some insight into how focussed learning promotes continual task performance in humans. Under focussed learning, participants learned the two category boundaries more accurately and with reduced mutual interference. This was revealed by a reduced slope of the psychometric function for the irrelevant task during F200 training, and by a psychophysical model which separately modelled errors arising from boundary inaccuracy and those incurred by generalised forms of forgetting. Without investigation of neural signals, it is challenging to draw strong inferences about how the task representations formed by focussed and interleaved learning may differ. For example, single-cell recordings in the macaque have suggested that after extensive (interleaved) training, neurons in prefrontal cortex form high-dimensional representations of task variables that are mixed across the neuronal population (dimension expansion), allowing efficient decoding of relevant actions via appropriate linear readout [23,24]. Mixed selectivity for distinct tasks will naturally similarly emerge in supervised neural networks after interleaved training, but not after focussed training, where instead representations oscillate periodically in line with the slowly-changing objectives. How, then, does focussed training promote robust task represenations in humans?

We suggest that one possibility is that focussed learning encourages humans to learn the structure of the stimulus space. Rather than encouraging dimension expansion, we propose that focussed learning encourages participants to learn a low-dimensional embedding of the stimulus space that maps cleanly onto the rules which could plausibly be useful for behaviour (in this case, leafy vs. branchy classification). Those participants with a prior bias to represent the trees according to their leafiness and branchines benefitted most from focussed learning, but only when the rules mapped onto the cardinal axes of the branch x leaf space. This benefit exceeeded that conferred during interleaved training, ruling out generalised explanations for this phenomenon. Our finding may relate to previous demonstrations that category learning benefits from “idealised” training exemplars which are unrepresentative of the test set but may promote understanding of the stimulus space [25,26]. Indeed, there are hints from the category learning literature that the benefits of interleaved training may be reduced or even reversed where there is low similarity among exemplars, rendering the diagnostic features hard to discover [27,28].

Finally, we harnessed these insights into human learning to conduct a proof-of-concept simulation using neural networks. Our goal here was not to achieve state-of-the-art performance in machine learning classification using our tree dataset. Indeed, as we show, ceiling performance on our task can easily be achieved with a standard convolutional neural network via interleaved training. Rather, we were interested in understanding the representations formed by the network during focussed training, which exhibits the temporal autocorrelation typical of naturalistic environments, and how these might be altered by exposure to unsupervised pretraining that encouraged the network to form appropriate embeddings of the two-dimensional tree space. We found that using a deep generative model as a feature extractor for the CNN partially guarded against catastrophic interference, and encouraged the preservation of representations of task A when later performing task B. One interpretation of this finding is that human continual learning is scaffolded by extensive unsupervised training that confers an understanding of the statistics of the world prior to any rule learning. However, we should note that this unsupervised structure learning alone is unlikely to be sufficient to explain the human capacity for continual learning over the lifespan. Indeed, it is very likely that other mechanisms, including weight-protection schemes that allow new learning to be allocated to synapses according to their importance for past learning, will also play a key role [2,29]. A promising avenue for future research may be to understand how structure learning and resource allocation schemes interact.

## Acknowledgements

This work was funded by an ERC Consolidator Grant (725937) to C.S., and by the Glyn Humphreys Memorial Scholarship to R.D.

## Author Contributions

C.S., T.F. and J.B. conceptualised the study, C.S. and T.F. designed the behavioural experiments, J.B. collected the behavioural data, T.F. designed and performed the Neural Network simulations, T.F. analysed the data, T.F., R.D., J.B., and H.N. provided new reagents/analytic tools, H.N. and C.S. provided expertise, C.S. and T.F. wrote the paper.

## Methods

### Participants

The study was approved by the University of Oxford Central University Research Ethics Committee (approval-no. R50750/RE001). Participants were adults who provided consent prior to taking part in the experiment in accordance with local ethics guidelines. We recruited a large cohort (n = 768) of participants via the online platform provided by Amazon Mechanical Turk with restrictions to the US (Exp.1) and India (Exp.2). We set a criterion of >55% accuracy at test for inclusion in the analysis, and continued to recruit participants until we reached at least n = 40 in each training group; attrition rates were high (but comparable with previous reports [28]), presumably because many participants were unused to conducting online tasks that involved uninstructed rule discovery. In total, we included 352 male and 231 female participants, with a mean age for included participants of 33.33 (range 19-55); ages did not differ reliably between groups. Final inclusion numbers and other details are detailed in the Table S1.

### Stimuli

Trees were generated with a custom fractal tree generator, written in Python 2.7 with the PIL package. The two feature dimensions were each varied parametrically in five discrete steps, spanning a two-dimensional space of leafiness x branchiness. We manipulated branchiness in terms of the branching angle, branching probability and the total number of recursive calls of the generator. Leafiness was defined as total number of leaves per branch, relative to branch length. Furthermore, positioning of individual leaves in front or behind the tree, leaf size, leaf rotation and individual branch colour tones were varied probabilistically. We created independent training and test datasets with of 100 trees each, and equal numbers of exemplars per combination of branchiness and leafiness. Thus, the same trees were shown for both tasks, to rule out low-level differences in saliency as influence on accuracy. Furthermore, different trees for training and test ensured that participants couldn’t rely on rote-learning the mappings between individual trees and responses. Each tree was shown eight times per task during training, and four times per task during the test phase. The data set was normed to equate difficulty along the two dimensions in a pilot study not reported here.

### Task and Procedure

Exp.1 and 2 were written in JavaScript with the Raphaël Vector Graphics Library and run in forced-fullscreen mode. We tested software compatibility with a variety of operating systems and web browsers before launching the task. All experiments consisted of a training phase (400 trials) and a test phase (200 trials). Self-paced breaks were allowed after 200 and 400 trials. We equated the number of presentations of each stimulus (leafy x branchy level). For Exp. 2, a similarity rating task was added before and after the main task (details below). At the beginning of the experiments, the participants received written instructions. They were asked to imagine that they owned two gardens, one in the north and one in the south. The goal was to learn via trial and error which type of trees grow best in each garden. Our instructions thus avoided alerting participants to the task-relevant dimensions (leafiness and branchiness).

In the training block, each trial began with the presentation of a centrally presented garden image (task context) for 500ms. Subsequently this image was blurred and a tree stimulus was presented for up to 2000ms. Responses were allowed 2500ms. As a reminder, the response assignments (e.g. keys for “plant” and “reject”) were displayed above the tree stimulus through the trial (counterbalanced). Feedback was displayed for 1500ms. Feedback on “plant” trials during training depended on the distance to the relevant boundary (+-50 for levels 1 and 5, +-25 for levels 2 and 4 and 0 for level 3). Reward was zero on reject trials. On plant trials, feedback was also accompanied by an animation (during which the garden was unblurred) that showed the tree grow or proportionally to the magnitude of the received reward. These timings were identical in test blocks, with the exception that no feedback was provided.

In Exp.1a and Exp.2a, the correct decision depended on leafiness for one garden and branchiness for the other (“cardinal” boundary). We fully counterbalanced the rule, so that equal number of participants received positive rewards for planting trees that were leafy/non-leafy and branchy/non-branchy. In Exp.1b and Exp. 2b, the decision boundary was rotated by 45 degrees, to align with the diagonal axes of the branch-x-leaf stimulus space. Thus, instead of learning the mapping from single feature dimensions onto rewards, participants had to learn the mapping of cardinal feature *combinations* onto rewards (“diagonal” boundary).

The test phase consisted of two trials in which the two contexts (gardens) were interleaved randomly over trials (100 instances of each). In Exp.1, we trained participants in 4 conditions, each involving 200 north and 200 south gardens in different order. In the interleaved condition, gardens were randomly interleaved over trials. In the F2, F20, and F200 conditions, gardens remained constant over 2, 20 or 200 trials (the first garden was selected at random). In Exp.2, we included only the F200 and interleaved training conditions.

### Arena task (Exp. 2)

In Exp.2a and 2b we additionally asked participants to rate the similarity among pools of 25 trees before and after the main experiment. We presented paricipants with a grey circular arena covering most of the screen area. This was populated by 25 trees (one per level of leafiness/branchiness) that were positioned in a random but non-overlapping configuration. Participants were asked to move the trees around by mouse drag-and-drop, arranging the trees so that more similar exemplars were close together and more dissimilar exemplars were further apart. The opacity of the selected tree was changed during the dragging process for usability purposes. This procedure was self-paced. Once all trees had been arranged and the participant was satisfied with the outcome, the next trial could be initiated by clicking on a designated button. Each trial involved a unique tree set. Participants performed 5 trials in total, taking an average of ~10 minutes.

### Analysis

To display learning curves, we plotted training accuracy in bins of 50 trials, and test accuracy averaged over the entire 200 trials. Accuracies were compared with ANOVAs. Other measures were compared with nonparametric tests, as violations of gaussianity were observed. For psychometric curves, for each participant we calculated p(plant) during the test phase for each of the 5 levels of the relevant and irrelevant dimensions, and fit a logistic function to the data. We compared the resulting parameters across the various cohorts using nonparametric statistics (ranksum test). For RSA analyses, we calculated p(plant) as a function of every level of leafiness x branchiness, and then computed an RDM expressing the dissimilarity (in accuracy) between each pair of leaf x branch level. We compared these to model-predicted RDMs that were generated to match a theoretically perfect observer (model 1) or an observer who learned the best possible single linear boundary through the 2D space and applied it to both tasks (model 2). In order to deal with partial correlation among these predictor matrices, they were orthogonalised using a recursive Gram-Schmidt approach. We computed Kendall’s Tau correlations between predicted and observed RDMs and compared these using a nonparametric approach (Wilcoxon Rank Sum Test).

Finally, we build a full psychophysical of the task. In a first step, we assumed that each stimulus was initially represented by a decision variable that was proportional its distance to a boundary in leaf x branch space. The boundary had signed angle φ with respect to the correct boundary. This value was converted to a choice probability via a logistic function with slope s, bias b and lapse parameter ε. We first ran a parameter recovery study to verify that the parameters were identifiable. The four parameters were fit to the human data via maximum likelihood estimation. We compared the best-fitting parameters across groups using nonparametric statistics.

### Influence of Priors

In Exp.2, for each participant and condition we calculated average pairwise distances between trees for each point in branch x leaf space, yielding an RDM of the same size as that described above (25 x 25). We then compressed this RDM to 2 dimension using multidimensional scaling. To compute the “grid” prior, for each participant we constructed a model RDM which exhibited perfect grid-like encoding of the two feature dimensions and correlated it with the empirical single-subject RDMs obtained from the arena task the participants enganged in prior to the main experiment. (Kendall taua rank correlation). We then repeated the analyses of test accuracy and the RDM analyses described above separately for participants with high and low griddiness prior, as indicated by a median split. We also conducted ANCOVA analysis on the Kendall’s Tau correlations with grid prior as a covariate of interest.

### Neural Network Simulations

All models were implemented in python 3.5 with tensorflow 1.1 and received RGB images as input. Details about the architectures and hyperparameter settings are reported in the supplementary methods. We generated training and test data sets with 50000 and 10000 trees respectively, using the same parameter settings as for the behavioural studies. In Exp 3, the network was trained in online mode (one sample per time) on 10000 trials per task, which were randomly sampled from the training data and superimposed onto the contextual cues (gardens). We trained the networks on focussed and interleaved curricula. Generalisation (test) performance on both tasks was assessed on independent data after the first and second half of the training session. For each combination of boundary and curriculum, 20 independent runs with different random weight initialisations were collected. In Exp 4, the β-VAE was trained until convergence on the training data with minibatches of n = 100. We then replaced the convolutional layers of the Exp. 3 network the the trained encoder and freezed the weights, such that training on the task took only place in the two fully-connected layers. Once again, we collected 20 independent runs per task.

To perform RSA, we fed independent sets of tree images with 10 exemplars per comination of branchiness and leafiness into the network and read out the (flattened) layer-wise activity patterns. RDMs were constructed from these activity patterns, yielding 50x50 RDMs (2 tasks x 25 trees) that captured between and within task dissimilarity structures for each layer. All subsequent analyses were identical to those reported for the behavioural studies. The model RDMs are described in detail in the supplementary material.

## Supplementary Material

Contains figures S1-S7, tables S1-S2 and supplementary methods.

**Supplementary Fig S1.**
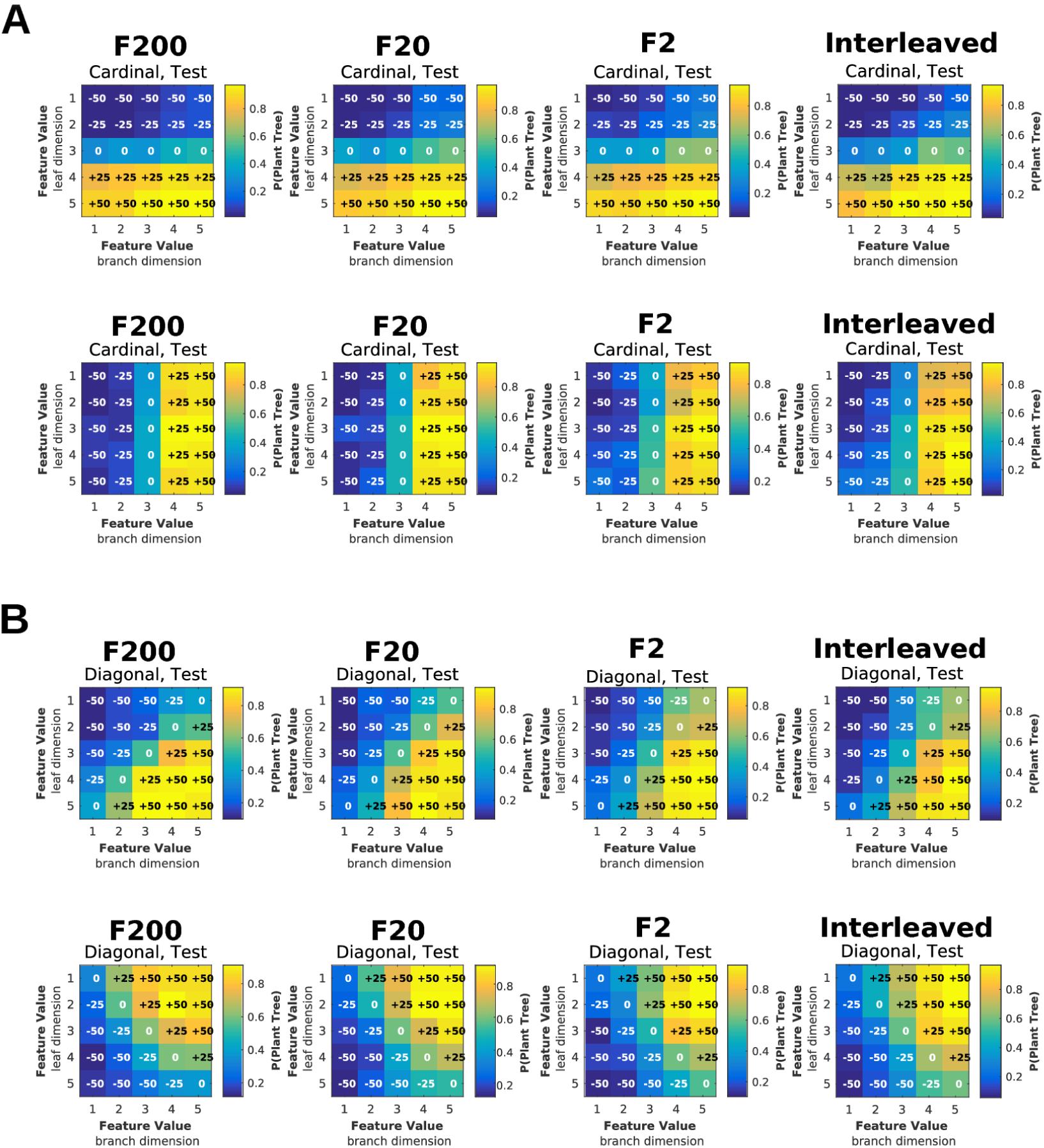
Exp 1. Group-level choice probability matrices. (A) Exp. 1a: Test-phase choice probability matrices for each task and group. One can nicely see a that patterns become more biased towards a combined boundary, the less the two tasks were temporally autocorrelated during training. Matrices were flipped and rotated on single subject level to account for counterbalancing of rewards signs across subjects. (B) Exp. 1b (diagonal): Test-phase choice probability matrices for each task and group. The patters resemble the ones described for. Whilst participants learned the boundaries very well under F200 training, interleaved or F2 training would lead to more mixed responses. In contrast to the cardinal boundary, there seem to be more random errors.

**Supplementary Fig S2.**
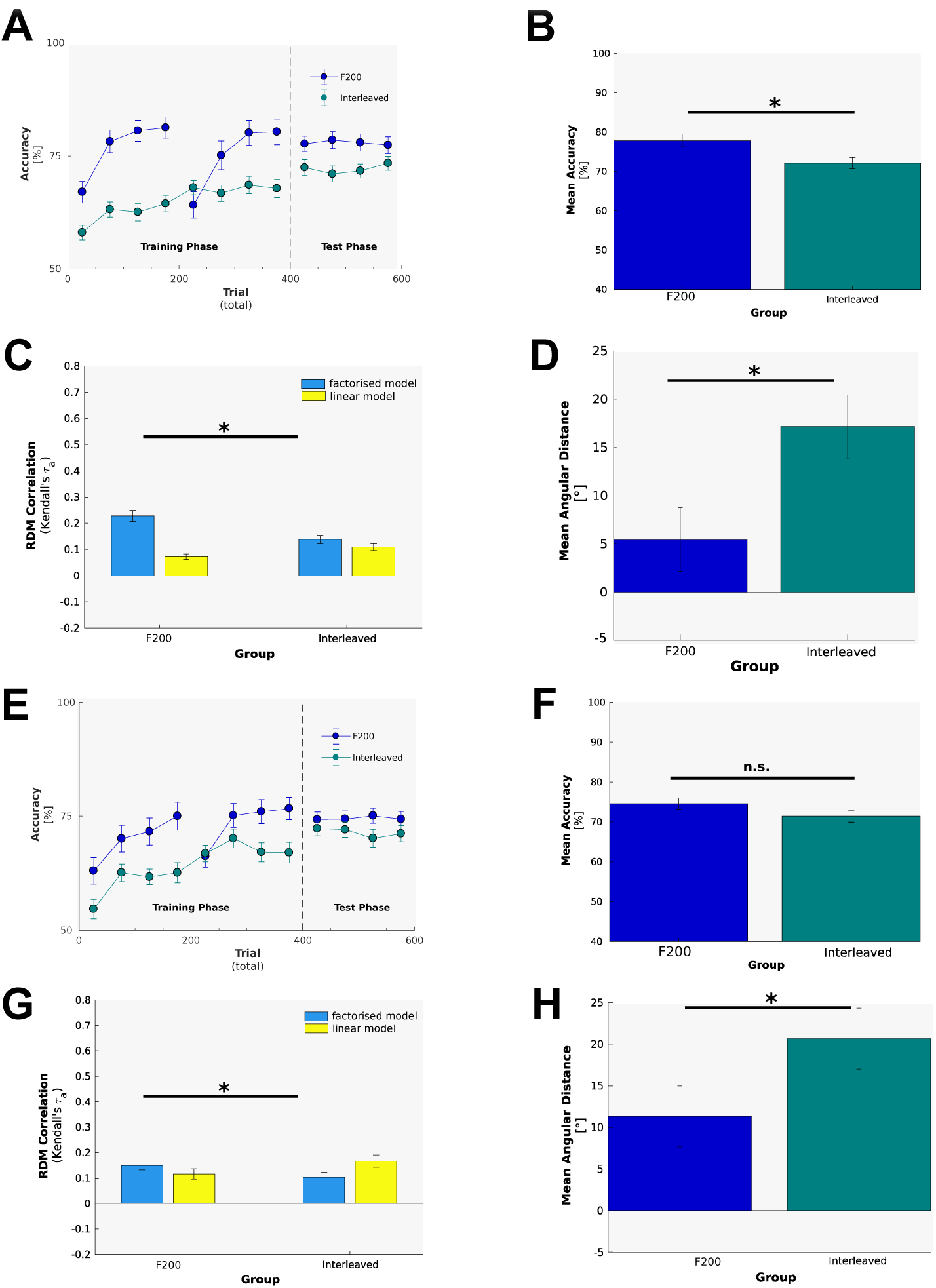
Exp 2. Replication of findings from Exp 1. (A) Exp. 2a, Learning curves. At the end of the training phase, both groups had reached a stable performance plateau. The F200 group learned fast and reached a higher training performance than the Interleaved group. This advantage continued during the test phase. (B) Exp. 2a. Mean test performance for F200 and Interleaved groups. Focussed training led to significantly higher test accuracy than interleaved training. (C) Exp2a. RDM Model correlations. There is stronger evidence for factorised representations after F200 than after interleaved training. Furthermore, interleaved training results in higher correlations with a linear model, than F200 training. (D) Exp. 2a, test phase decision boundary bias, estimated by psychophysical model. Interleaved training resulted in a significantly stronger difference between estimated decision boundary and true category boundary, than F200 training. (E) Exp. 2b, learning curves. While the F200 group learned faster and reached a higher terminal training performance than the interleaved group, the differences in the test phase were rather minute. (F) Exp. 2b, test phase mean accuracy. As for Exp. 1b, there was no significant difference in test performance between F200 and interleaved training on diagonal boundaries. (G) Exp. 2b, RDM model correlations. The factorised model explained the F200 group data – but not the interleaved group data – better than the linear model. (H) Exp 2b, decision boundary bias. Participants in the interleaved group had a significantly higher bias in their estimated decision boundaries than participants in the F200 group.

**Supplementary Fig S3.**
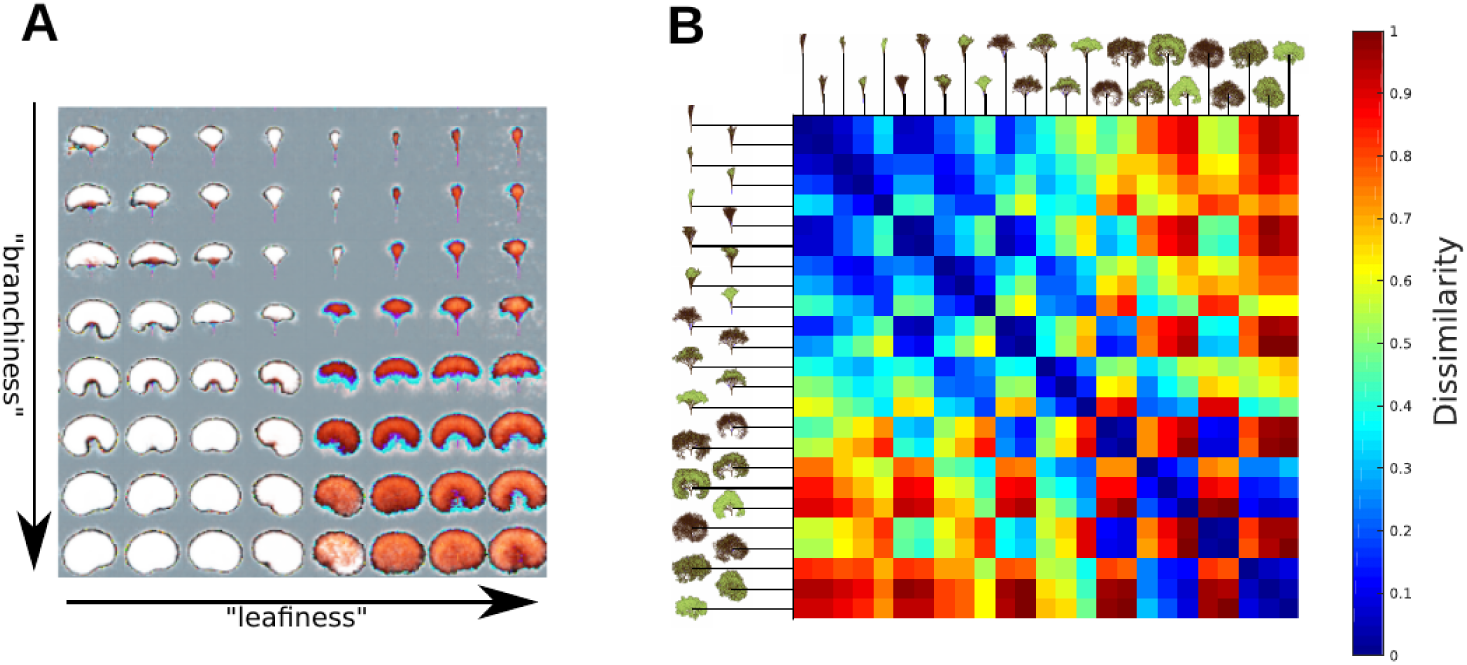
Exp 4a, Latent Space Traversal and Encoder RDM. (A) Exp. 4a. Latent space traversal. Here, we traversed the two-dimensional latent space of the β-VAE and generated tree images via sampling from the latent distribution and feed-forward passes through the decoder. The generated tree images were then placed at their corresponding positions in the latent space. Interestingly, for this particular choice of beta (=50), the latent variable layer developed disentangled and interpretable factors that seem to represented “leafiness” and “branchiness”. (B) Exp 4b. RDM obtained from the activity patterns in the last encoder layer of the trained β-VAE (beta=50). Interestingly, even this layer seemed to encode both dimensions (leafiness and branchiness) as indicated by the checkerboard pattern (=leafiness) and higher similarity along the main diagonal (=branchiness). We interpreted this as evidence that the even the β-VAE encoder can be used as feature extractor for the main task.

**Supplementary Fig S4.**
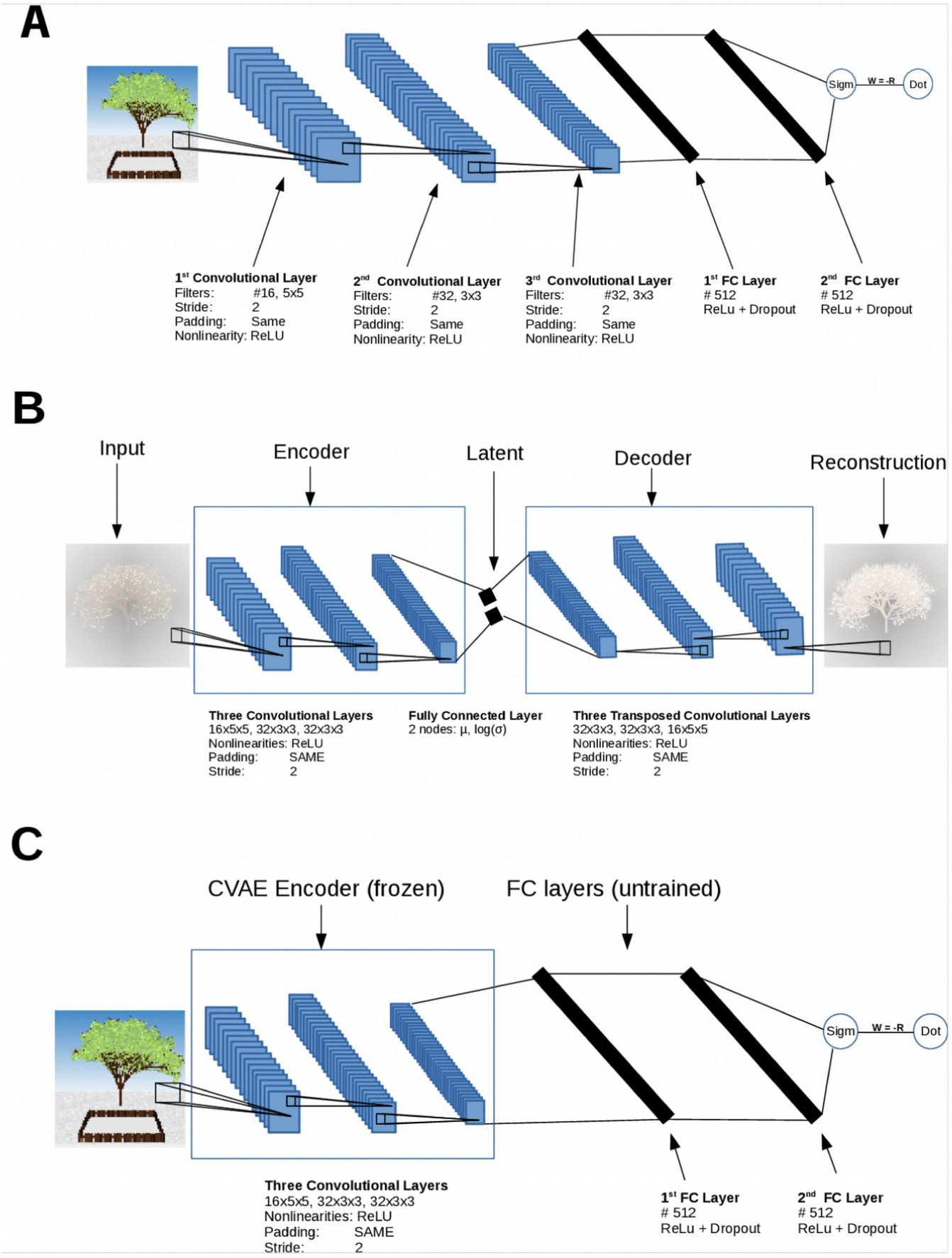
Neural Network Architectures. (A) Exp 3. Model Architecture. The agent consisted of three convolutional layers(16x5x5,32x3x3,32x3x3, padding=SAME, nonlinearity=ReLU, stride=2) and two fully-connected layers (512x1,512x1, nonlinearity=ReLU, dropout=0.5), followed by a one-dimensional output unit (nonlinearity = Sigmoid). Weights were initialised with He-Initialisation, and biases set to 0.01. (B) Exp 4a. Model Architecture. The encoder consistent of three convolutional layers which were identical to the first three layers of the network depicted in. We flattened the output of the encoder and fed it into the two-dimensional latent layer, which was separated into a node for the mean paramenter and a node for the logsima parameter of the latent distribution. These parameters were combined with a sample from a zero mean, isotropic variance gaussian distribution in the subsequent reparametrisation layer. The decoder consisted of three layers with transposed convolutions, arranged in reverse order of the encoder layers. All layers in the encoder and the first two decoder layers passed their outputs through ReLU nonlinearities. No nonlinearities were applied to the latent layer and the final output of the decoder was passed through a hyperbolic tangent, as the input image could have values below zero due to normalisation, and the optimisation aim of the VAE was to resemble the input in its output as close as possible. The weights were initialized using He-Initialization and all biases were initially set to 0.001. (C) We selected the encoder of the model with the highest gridiness score in Exp. 4a and used it to replace the convolutional layers of the agent from Exp. 3. Note that architecture is exactly the same, but weights now instead of being randomly initialised encode an efficient representation of the stimulus space before any training on the trees task has taken place. Next, we freezed the weights of the convolutional layers to ensure that the representation remains persistent and serves as feature extractor throughout training.

**Supplementary Fig S5.**
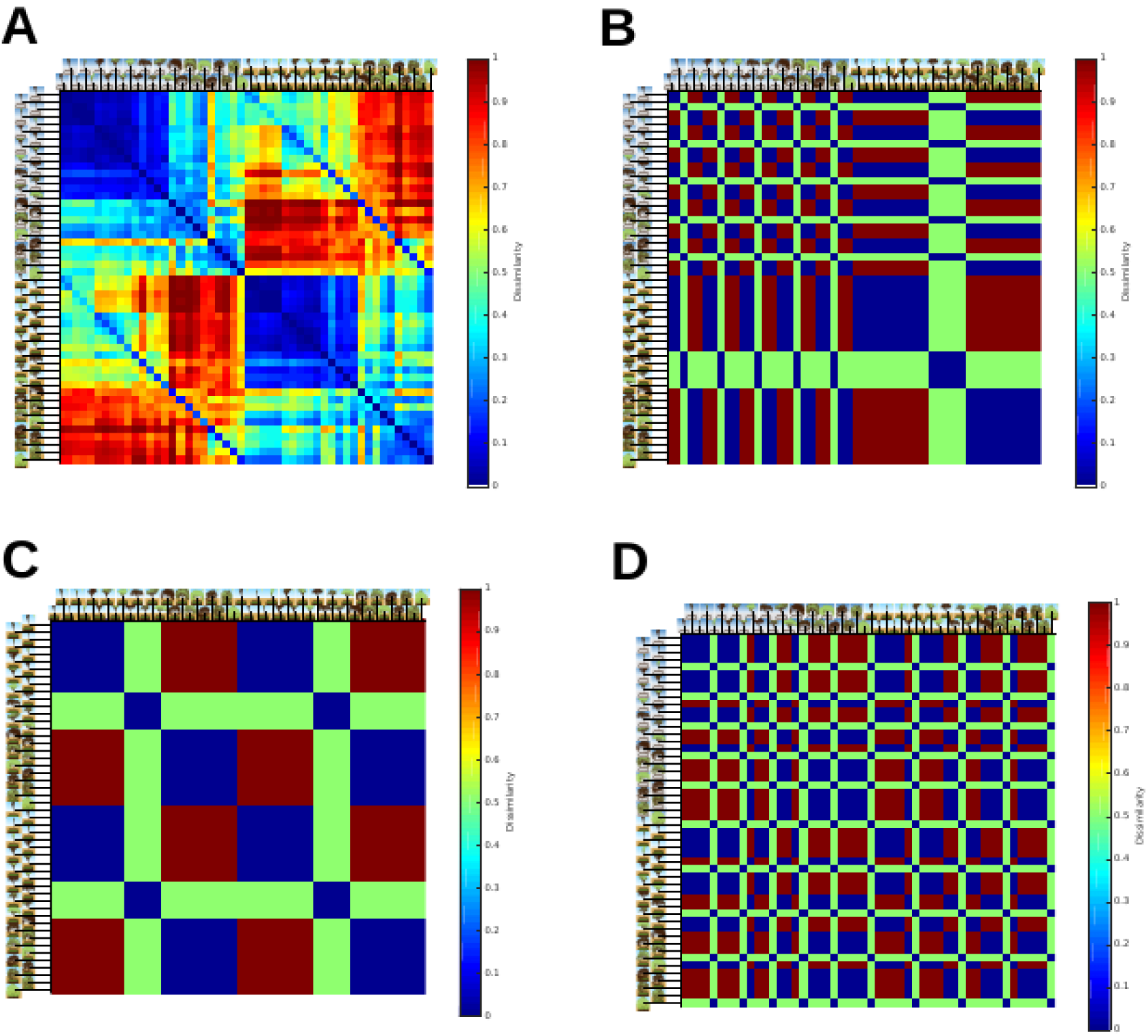
Exp3,4. Example of Model RDMs. (A) Pixel Space dissimilarity model. Here, we computed the dissimilarity between vectorised RGB images for all 50 possible combinations of leafiness,branchiness and task. (B) Factorised Model. Cardinal boundaries. This model RDM assumed that the network had learned perfect binary representations of both categories for each task (compare). (C) Interference Model RDM. Cardinal boundaries. If the network had been trained first on the north and subsequently on the south task, we expected catastrophic interference. This model RDM assumes that the first task is treated as if it was the second task. (D) Linear Boundary Model. Cardinal boundaries. In the behavioural experiments, we tested the hypothesis that participants learn only one single, *linear* boundary for both tasks, which is positioned in trees space such that reward is maximised in both gardens. This RDM illustrates how representational geometries would look like in such a model.

**Fig S6.**
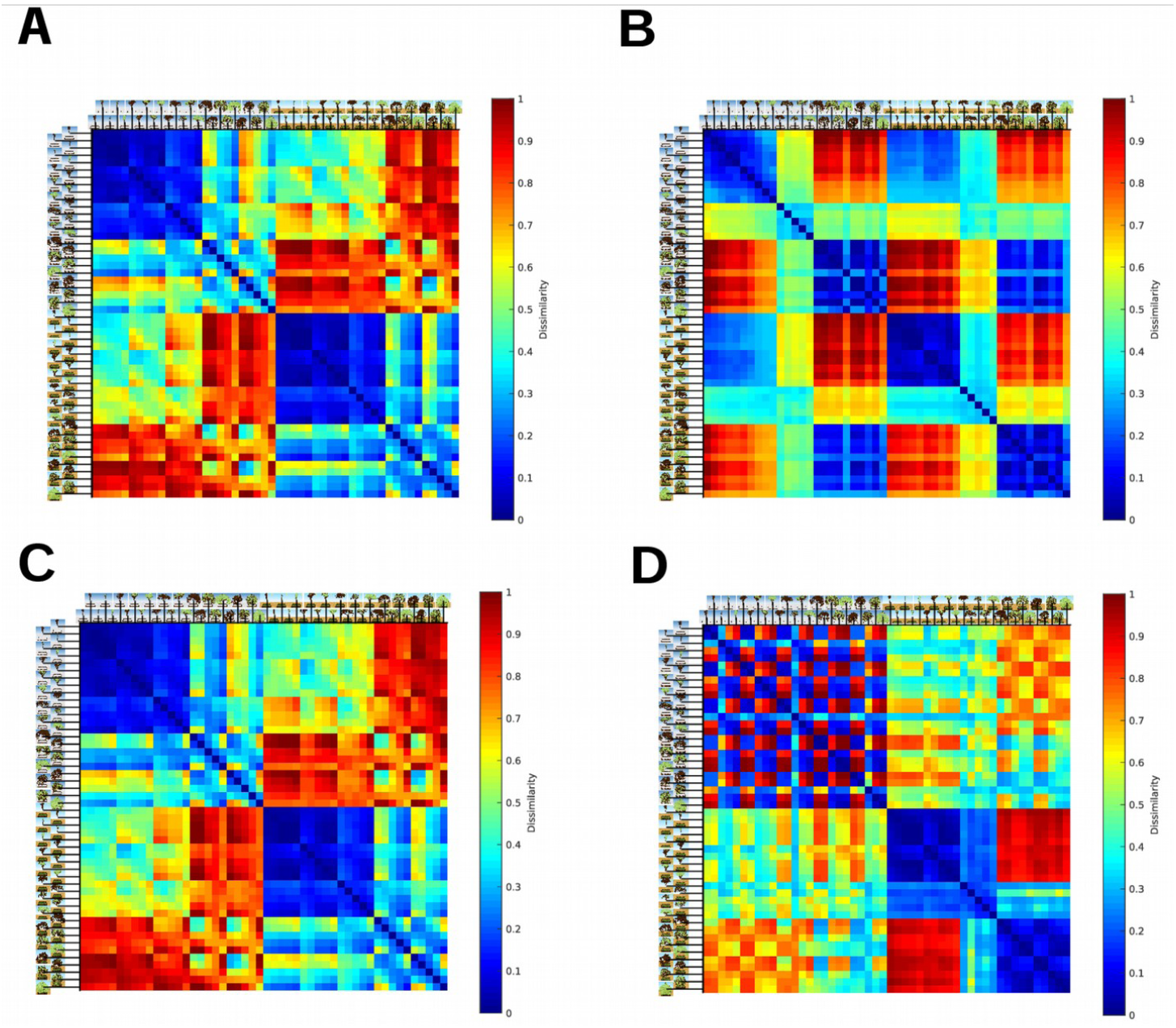
Exp 3a, Example of Layer RDMs. (A) Exp 3a, focussed training. RDM constructed from the activity patterns in the first convolutional layer during the final test phase. (B) Exp 3a, focussed training. RDM constructed from the test-phase activity patterns in the FC2 layer. As expected, the network responds to the first task (top left quadrant) as if it was the second task (bottom right quadrant). (C) Exp. 3b, interleaved training. RDM obtained from the first convolutional layer during the last test phase. As under focussed training, mostly differences in pixel value patterns is encoded. (D) Exp 3b, interleaved training. RDM obtained from the FC2 layer. In contrast to focussed training (Fig S6b), the second FC layer exhibits clear separation of representational geometries for the north and south task. In other words, interleaved training allowed the network to learn the boundaries for each of the two tasks, which is reflected in FC2-layer responses to branchiness in the south and leafiness in the north task.

**Supplementary Table S1.**
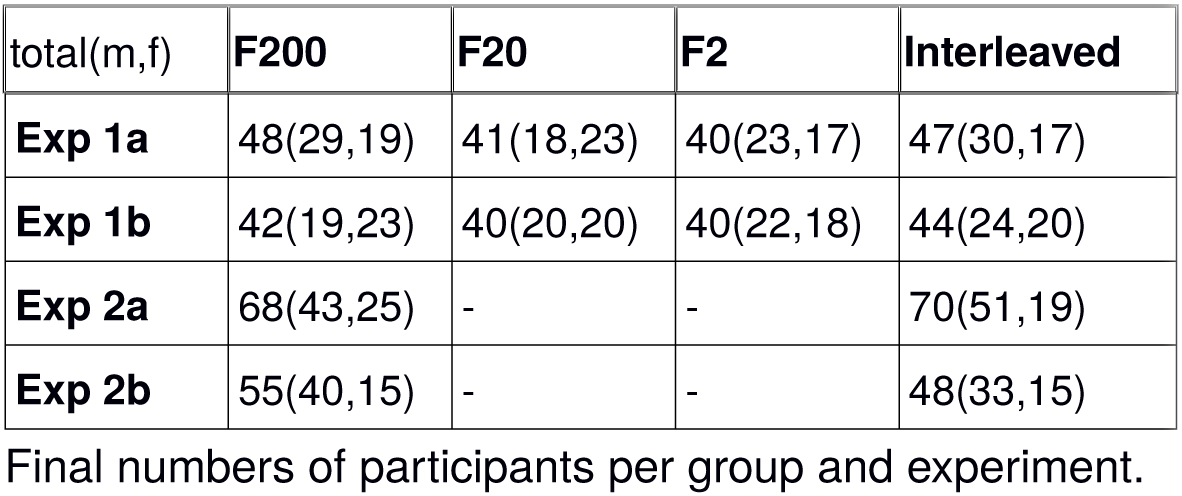

| total(m,f) | F200 | F20 | F2 | Interleaved |
| --- | --- | --- | --- | --- |
| **Exp 1a** | 48(29,19) | 41(18,23) | 40(23,17) | 47(30,17) |
| **Exp 1b** | 42(19,23) | 40(20,20) | 40(22,18) | 44(24,20) |
| **Exp 2a** | 68(43,25) | - | - | 70(51,19) |
| **Exp 2b** | 55(40,15) | - | - | 48(33,15) |

Final numbers of participants per group and experiment.

**Supplementary Table S2.**

|  | F200 | F20 | F2 | Interleaved |
| --- | --- | --- | --- | --- |
| **Exp 1a** | 34.38 | 34.22 | 35.85 | 35 |
| **Exp 1b** | 34 | 37.1 | 34.5 | 36.82 |
| **Exp 2a** | 31.15 | - | - | 30.71 |
| **Exp 2b** | 32.89 | - | - | 29.38 |

Mean age of participants per group and experiment.

## Supplementary Procedures

### Exp 3,4: Neural Network Simulations

All models were implemented in python 3.5 with tensorflow 1.0 and trained in GPU mode on an Nvidia Tesla K80 GPU.

### Exp 3: CNN Simulations of Trees task without priors

In this experiment, we replicated the trees study with a convolutional neural network (CNN), which learned solely from pixels and a reward signal. Goal was to compare human performance and choice biases with the effects of blocked vs interleaved learning on its hidden layer representations and test performance, using the same task. The network architecture is described in **Fig. S4a**.

#### Training

We generated 50000 training and 10000 test tree images using exactly the same settings as for the behavioural experiments. The tree images were downscaled to 96x96x3 and saved as .png RGB images.

Training occurred online with one sample per time. Each training trial consisted of a composite input image, consisting of the garden and a superimposed tree. This image was fed through the network, to create a choice probability. We used the negative reward, multiplied by the choice probability, as custom loss function. The agent was trained via stochastic gradient descent (SGD) with the Adam optimiser and a learning rate of 1e-4.

#### Task Design and Procedure

We compared the effects of blocked and interleaved training. In the blocked curriculum, the agent was first trained on 10000 trials from one task, and then on 10000 trials from the second task. In the interleaved group, the agent was trained on 20000 randomly shuffled trials from both tasks. In both cases, we evaluated the performance on 10000 previously unseen test trials per task. The training data was sampled from the set of 50000 trees, keeping the number of trees per combination of leafiness and branchiness consistent across tasks and runs. The test phases occurred after half of the training trials were fed through the network, and at the end of the training phase. During these evaluation phases, we recorded the activations in the convolutional, fully-connected and output layers on each trial.

We counterbalanced task order and reward assignments in the same way as for the behavioural experiments. In total, 40 independent runs with random weight initialisations per combination of boundary (cardinal or diagonal) and curriculum (blocked or interleaved) were collected.

#### Analysis

To evaluate performance, we binned the training phase into bins of 100 trials, and computed the percentage of correct choices for each bin. This was carried out on independent runs, before we computed the group means for each training curriculum. For the test phase, we calculated the percentage of correct decision across all trials, for both tasks individually and across tasks, before averaging across runs. In both cases, we excluded category boundary trials, where no correct decision was possible.

Furthermore, we constructed Representational Dissimilarity Matrices (RDMs) from the individual layers, based on their responses to test data stimuli. The procedure was almost identical to our RSA on the behavioural data. However, instead of having just one value (e.g. choice probability) per stimulus, we obtained now response matrices of size 50xn with n being the size of the (flattened) layer activations. The dissimilarity between activity patterns for pairs of stimuli was calculated using the correlation distance measure (1-correlation), yielding one 50x50 RDM per layer. Once again, we correlated these RDMs with model RDMs (**Fig. S5**), capturing pixel dissimilarity, factorised task encodings, catastrophic interference (e.g. encoding of task 1 as if it was task 2) or a shared linear boundary.

### Exp 4a: Unsupervised Learning of Disentangled Representations

Aim of this experiment was to simulate the unsupervised learning of the stimulus space and emergence of disentangled visual concepts (of branchinesss and leafiness). We attempted to show that the most efficient compression of the data would actually correspond to the structure of the trees space. The network architecture is illustrated in **Fig. S4b**.

### Training Procedure

The training data set consisted of 50000 trees stimuli, with equal numbers of examples per combination of branchiness and leafiness. Another set of 10000 trees stimuli, equally balanced, was generated and served as test set. All tree stimuli were scaled to 96x96x3 pixels. We normalised the data separately for each colour channel by subtracting the mean and dividing by the standard deviation. The network was trained with minibatches of 128 training stimuli and training was stopped after 20 epochs on the entire training data, as we noted that the network would otherwise overfit to the training data. We evaluated the network’s performance after each epoch with a full pass of the test data. Furthermore, after each trianing epoch, a latent space traverse (see below) was performed and the encoder layer outputs for two full 5x5 sets of tree stimuli were stored for later Representational Similarity Analysis.

Training was repeated for different values of beta, ranging from 1 to 100. We found a beta of 50 to yield the best disentanglement (see below).

### Analysis

For the latent space traversal, we fed linearily spaced values for the mean parameters, ranging from -2 to +2 along each of the two latent dimensions into the network and generated the tree via a full pass through the decoder. For each of these value pairs, we placed the generated image at the corresponding (x,y)-position in latent space, to obtain a visualisation of the traverse. In doing so, we could qualitatively assess the degree to which each of the two latent dimensions captures unique variation along the relevant feature dimensions.

Furthermore, we constructed 25x25 RDMs out of the activations recorded from the encoder layer, using the same methodology as previously described. These RDMs could then be correlated with a conceptual model RDM which exhibited a structure one would obtain if the stimuli were represented in a perfectly linear grid, according to their dissimilarity in branchiness and leafiness. Calculating these RDM model corelations (Kendall taua) yielded one “grid score” per beta parameter. This allowed us to identify the value for beta which resulted in the most grid-like representations in the last encoder layer.

### Exp 4b: CNN Simulation of Trees task with priors

The previous experiment identified a parametrisation for the VAE which allowed us to model our participant’s “awareness” of the stimulus space structure. That is, the model learned a compressed representation which arranged the stimuli according to their variation in leafiness and branchiness. As **Exp 2** revealed that participants with a high gridiness score, e.g. with a strong awareness of the stimulus space, performed better in the blocked curriculum, compared to subjects with a lower “prior”, we could now ask if providing a neural network with such representations would boost its continual task performance.

#### Training, Procedure and Analysis

The training parameters and task design were identical to the ones described for Exp. 3. We collected the same number of runs per group and recored the hidden layer activity on independent test data. Likewise, the same analyses were performed, namely calculation of learning curves, test performance and RSA on the layer-wise activity patterns.

